# Smc5/6-antagonism by HBx is an evolutionary-conserved function of hepatitis B virus infection in mammals

**DOI:** 10.1101/202671

**Authors:** Fabien Filleton, Fabien Abdul, Laetitia Gerossier, Alexia Paturel, Janet Hall, Michel Strubin, Lucie Etienne

**Author notes:** Contributed equally to this work (Fabien Filleton, Fabien Abdul). Correspondence: Lucie Etienne, CIRI, ENS-Lyon, 46 Allée d’Italie, 69364 Lyon, France.

## Abstract

Infection with Hepatitis B virus (HBV) is a major cause of liver disease and cancer in humans. HBVs (family *Hepadnaviridae*) have been associated with mammals for millions of years. Recently, the Smc5/6 complex, known for its essential housekeeping functions in genome maintenance, was identified as an antiviral restriction factor of human HBV. The virus has however developed a counteraction mechanism by degrading the complex *via* its regulatory HBx protein. Whether the antiviral activity of the Smc5/6 complex against hepadnaviruses is an important and evolutionary-conserved function is unknown. Here, we used a combined evolutionary and functional approach to address this question. We first performed phylogenetic and positive selection analyses of the six Smc5/6 complex subunits and found that they have been highly conserved in primates and mammals. Yet, the Smc6 subunit showed marks of adaptive evolution, potentially reminiscent of virus-host “arms-race” We then functionally tested the HBx from six very divergent hepadnaviruses now naturally infecting primates, rodents, and bats. Despite little sequence homology, we demonstrate that these HBx efficiently degraded mammalian Smc5/6 complexes, independently of the host species and of the sites under positive selection. Importantly, all also rescued the replication of an HBx-deficient HBV in primary human hepatocytes. These findings point to an evolutionary-conserved requirement for Smc5/6 inactivation by HBx, showing that the Smc5/6 antiviral activity has been an important defense mechanism against hepadnaviruses in mammals. Interestingly, Smc5/6 may further be a restriction factor of other yet unidentified viruses that have driven some of its adaptation.

**Importance:** Infection with hepatitis B virus (HBV) led to 887000 human deaths in 2015. HBV has been co-evolving with mammals for millions of years. Recently, the Smc5/6 complex, known for its essential housekeeping functions, was identified as a restriction factor of human HBV antagonized by the regulatory HBx protein. Here, we address whether the antiviral activity of Smc5/6 is an important evolutionary-conserved function. We found that all six subunits of Smc5/6 have been conserved in primates with only Smc6 showing signatures of “evolutionary arms-race” Using evolutionary-guided functional assays that include infections of primary human hepatocytes, we demonstrate that HBx from very divergent mammalian HBVs could all efficiently antagonize Smc5/6, independently of the host species and sites under positive selection. These findings show that the Smc5/6 antiviral activity against HBV is an important function in mammals. It also raises the intriguing possibility that Smc5/6 restricts other, yet unidentified viruses.

## Introduction

Hepatitis B virus (HBV) infects more than 250 million people worldwide and is a leading cause of chronic hepatitis and liver cancer in humans (World Health Organization). HBV is a member of the *Hepadnaviridae* family of DNA viruses, which have co-evolved with their host species for millions of years (1-3). Today, hepadnaviruses are found to naturally infect, in a species-specific manner, mammals, birds, as well as fish and amphibians. In mammals, HBVs are present in rodents, bats, and several primates including human, chimpanzee, gibbon, orangutan, and the New World wooly monkey. Mammalian hepadnaviruses (orthohepadnaviruses) all contain a gene encoding a small regulatory protein, HBx, that is thought to have arisen *de novo* in the orthohepadnavirus lineage (1). HBx has long been known to play a central role in HBV replication and pathogenesis (4-6) and has recently been shown to have a key role in promoting HBV transcription by antagonizing the restriction function of the infected cell’s Structural Maintenance of Chromosome (SMC) Smc5/6 complex (7, 8). However, whether this property has been conserved among the HBx-containing hepadnaviruses is unknown.

The Smc5/6 complex is, together with cohesin and condensin, one of the three SMC complexes found in eukaryotes (9, 10). As for the other SMC complexes, the core of the Smc5/6 complex is formed by a heterodimer of two SMC proteins, Smc5 and Smc6 (11), which associate with four additional subunits known as non-SMC elements (Nsmce1-4) (Figure 1A). These SMC complexes all have essential housekeeping functions, playing fundamental roles in chromosome replication, segregation and repair (reviewed in (10). Condensin controls chromosome condensation during mitosis and cohesin maintains cohesion between the newly replicated sister chromatids. However, the role of the Smc5/6 complex is less well understood. It has reported functions in DNA replication and repair, but its exact mode of action remains elusive (12-16).

**Figure 1.**
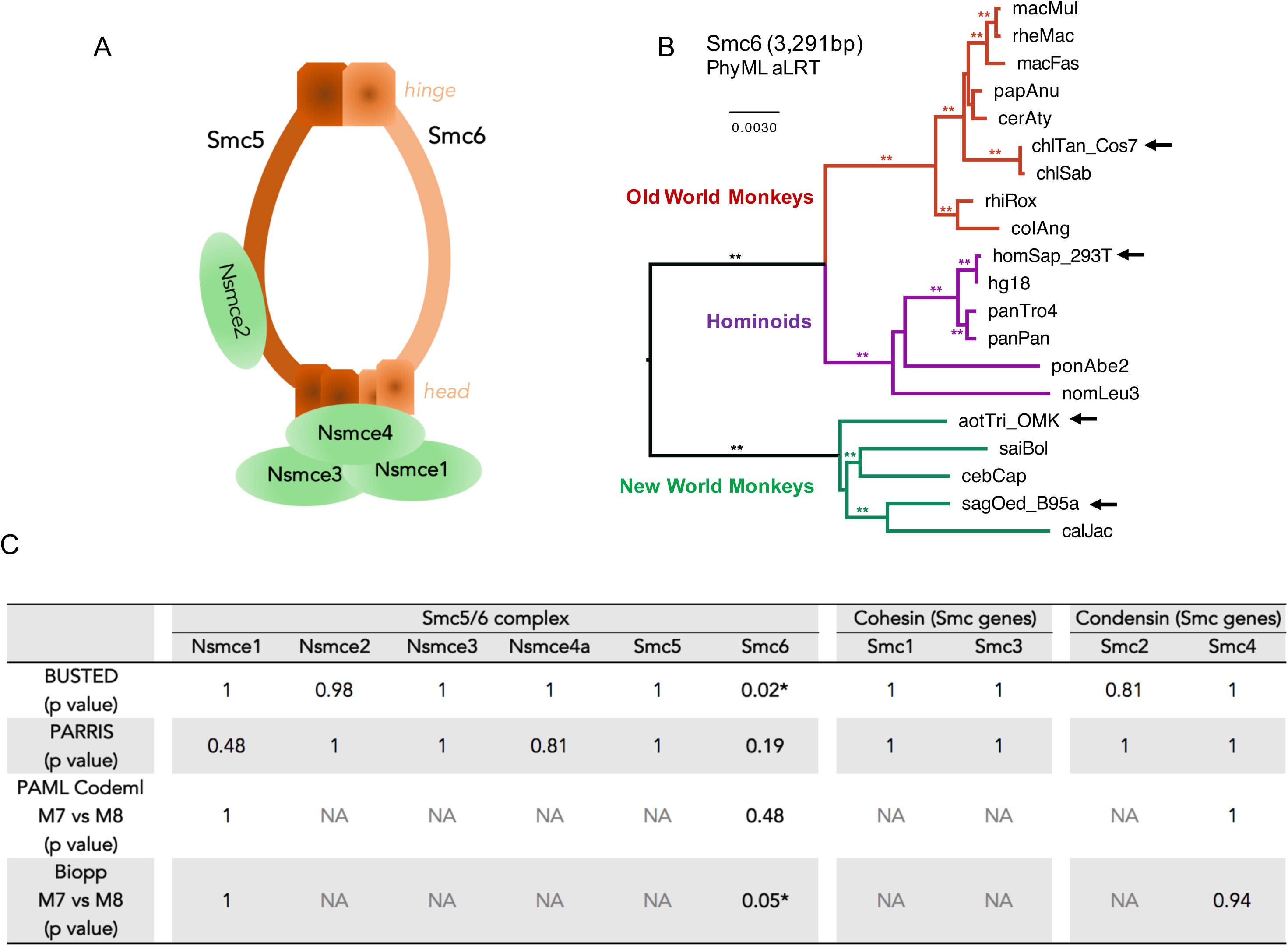
Smc6 is the least conserved subunit of the Smc5/6 complex in primates. A, Architecture of the Smc5/6 complex. The complex is made of two core subunits (Smc5 and Smc6) and four non-SMC elements (Nscme1-4). B, Phylogenetic analysis of primate *Smc6* genes. Sequences were aligned with MUSCLE and phylogeny was performed with PhyML and an HKY+I+G model with aLRT as statistical support (**, aLRT>0.8). Newly sequenced genes (arrow) are indicated. The newly sequenced *Smc6* from *Chlorocebus pygerythrus* (chlPyg_Vero cells) is not represented because the nucleotide sequence is identical to the retrieved *Chlorocebus sabaeus* (chlSab) sequence of *Smc6*. Alignments and phylogenies of the ten SMC analyzed genes are available in Supplementary Dataset S1. C, Positive selection analysis of the indicated genes during primate evolution. Shown are the p-values obtained using four different methods (BUSTED, PARRIS, PAML Codeml, and Bio++; see Methods). The p-values of the maximum-likelihood tests indicate whether the model that allows positive selection better fits the data (*: statistically significant). NA, results are not available because convergence was not obtained for these genes and/or analyses (see Methods).

In addition to its essential cellular activities, a novel function of human Smc5/6 complex as an HBV restriction factor has been recently dissected: in the absence of HBx, the Smc5/6 complex binds to the HBV episomal DNA genome and inhibits viral transcription (7, 8, 17). Human HBx antagonizes this effect by high-jacking the host DDB1-containing E3 ubiquitin ligase complex to target the Smc5/6 complex for ubiquitin-mediated degradation, thereby enabling productive HBV gene expression (7).

Most genes encoding for antiviral restriction factors have been engaged in an “evolutionary arms-race” with the viruses they inhibit (18, 19). Indeed, during long-term coevolution, pathogenic viruses and their hosts are constantly under the selective pressure of the other for survival. As a result, host restriction factors evolve rapidly and display signatures of positive (diversifying) selection. These signatures can be identified by analyzing the codon sequences of orthologous genes from a large number of related species. At virus-host interaction sites, one can witness adaptive changes, including frequent amino acid changes (where a higher non-synonymous substitution rate, dN, than the synonymous rate, dS, is indicative of positive selection), and insertion/deletions (indels) or splicing variants as ways to modify the virus-host interface and to escape from viral antagonists (19-24).

To assess whether the antiviral function of the Smc5/6 complex has been evolutionarily important, we performed phylogenetic and evolutionary analyses of virus and host proteins in combination with functional assays. We found that all six subunits of the Smc5/6 complex have been highly conserved in primate evolution, with only Smc6 showing signatures of an “evolutionary arms-race”. Because orthohepadnaviruses have diverged millions of years ago and their HBx have very little sequence homologies, we then investigated the Smc5/6-antagonism capacity of HBx from six divergent orthohepadnaviruses from primates, rodents and bats. We found that all orthohepadnavirus HBx are efficient at counteracting the Smc5/6 complex, independently of the host species or the variations at sites under adaptive evolution, and including in infections of human primary liver cells. Therefore, Smc5/6-antagonism is a strict requirement for the establishment of mammalian HBV infection, showing that the Smc5/6 complex has been an important immune defense against hepadnaviruses in mammals. Our findings also raise the intriguing possibility that the Smc5/6 complex may restrict other, yet unidentified pathogenic viruses.

## Results

### Overall evolutionary conservation of the Smc5/6 complex in primates

To trace the evolutionary history of the Smc5/6 complex, we compared the sequence of its six subunits in primates (Figure 1A). As a comparison, we also analyzed the evolutionary history of all the other primate SMC genes. These include the *Smc1* and *Smc3* genes, which encode the core cohesin subunits, and the *Smc2* and *Smc4* genes encoding the condensin core subunits. The sequences of these genes were retrieved from publically available datasets (Table 1, Dataset S1). To perform more robust phylogenetic and selection analyses, we obtained additional primate species sequences using RT-PCR approaches (Table 1, Figure 1B; see Methods). Overall, we included up to 20 simian primate species in our positive selection analyses to span 40 million years of evolution (25, 26). We found that the synteny of the genes was conserved during primate evolution, although some subunits had duplicated pseudogenes in a few primate species (Figure S1). Amongst the core SMC proteins, the most conserved are the cohesin Smc1 and Smc3 subunits, which share 100% and 99.9% pairwise identity at the amino acid level in a simian primate alignment, respectively (Dataset S1). Smc6 was the least conserved SMC protein with 97.4% pairwise identity in simian primates. Using the Genetic Algorithm for Recombination Detection (GARD) on the complete set of genes (27), evidence of recombination (GARD, p<0.05) was only found for *Nsmce3,* and therefore subsequent phylogenetic and selection analyses were performed on both the whole *Nsmce3* gene and the two identified *Nsmce3* gene fragments (from 1-246 bp and 247-912 bp). Phylogenetic analysis of the ten genes showed that the gene trees derived from nucleotide alignments were largely in accordance with the accepted species tree from Perelman and colleagues (Figure 1B, and Dataset S1) (26).

**Table 1.**
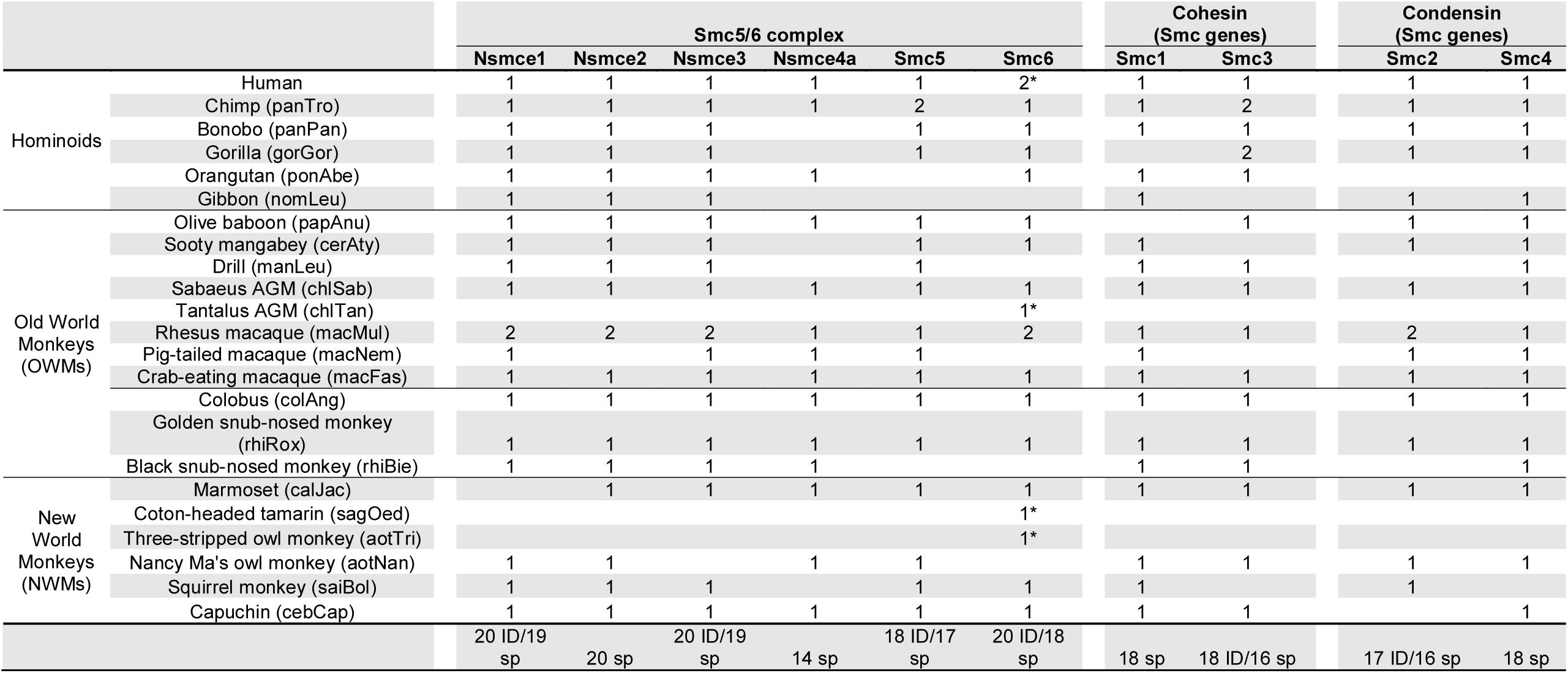
**Primate species and sequences that we included in the evolutionary analyses of the SMC complexes.** The asterisks indicate the sequences newly obtained in this study. The total number of primate species (sp) and individuals (ID) included in each gene analysis is indicated at the bottom. The abbreviated species names are found in parentheses (See Methods for details).

To assess whether the Smc5/6 complex and the *Smc1-4* genes have experienced diversifying selection during primate evolution, we performed four types of positive selection analyses. Firstly, we used the BUSTED method that tests whether a gene has experienced positive selection on at least one site or one branch during evolution (28). Of the six Smc5/6 complex subunits and the four *SMC* genes from the cohesin and condensin complexes, only one, *Smc6*, showed gene-wide evidence of episodic positive selection (BUSTED, p<0.05, Figure 1C). These findings were confirmed using the PARRIS method that also detects evidence of positive selection using a codon alignment (29), although the level of significance was not reached for the *Smc6* gene (p=0.19; Figure 1C). Thirdly, we ran the Codeml program from the PAML package (30) on our codon alignments to compare two models: one that allows positive selection at certain sites (M8, alternative hypothesis) and one that disallows positive selection (M7, null hypothesis). We then performed a likelihood ratio test (LRT) to examine which of the two models fitted better our data. Overall, there was no convergence for seven of the ten genes because they were too conserved (see Methods for details) and no evidence of significant positive selection was found for the remaining genes (Figure 1C). Finally, we used the Bio++ package from Guéguen *et al.* (31), which has two main advantages over PAML to similarly test for evidence of positive selection (M7 *vs.* M8 as implemented in Bio++). First, the DNA substitution models use non-stationary matrices, which allow nucleotide composition to change over time, and therefore improve the fitting to real data (32). Second, the codon frequency better fits biological assumptions, as compared to PAML (31). Using Bio++, we obtained higher values of likelihood and, for *Smc6,* a model allowing positive selection was favored over a model that disallows positive selection (Bio++ M7 *vs.* M8, p=0.05; Figure 1C). Overall, these studies show that the *SMC* genes for cohesin and condensin, as well as the six Smc5/6 complex subunits, have been highly conserved during primate evolution. This is similar to what has been described for the global evolution of SMC proteins in eukaryotes (9) and is consistent with their essential cellular housekeeping functions. However, it is in contrast to most other known antiviral restriction factors that have strongly evolved under positive selection in primates (18, 21). The only exception is the *Smc6* gene for which two positive selection analyses found evidence of adaptive evolution.

### Evidence of site-specific adaptive evolution in primate Smc6

It is formally possible that most of the Smc6 protein has been highly conserved due to its housekeeping function, whilst just a few sites have been engaged in a virus-host interaction. Indeed, previous studies have shown that essential cellular proteins that are usurped by viruses for their replication have evolved under a strong purifying selection background with only the virus-host interaction sites showing rapid evolution (33-37). To determine if this was the case for *Smc6*, we characterized in more detail its evolutionary history during primate evolution. Using the BranchSite-REL algorithm from HYPHY (38), we found that positive selection has occurred on a proportion of branches of the primate *Smc6* phylogeny (episodic positive selection; BS-REL, p<0.05). To look more specifically for site-specific positive selection, we used MEME and FUBAR, which are the most accepted methods from HYPHY, as well as the posterior probabilities at codon sites in PAML and Bio++ (M8 model) (Figure 2A). We confirmed that most codons in Smc6 have been extremely conserved, with over 90% having a non-synonymous to synonymous nucleotide (dN/dS) substitution ratio lower than 1 (Figure 2B). However, a few sites were identified as having evolved under significant positive selection by one or several methods, consistent with site-specific positive selection in the primate *Smc6* gene (Figure 2).

**Figure 2.**
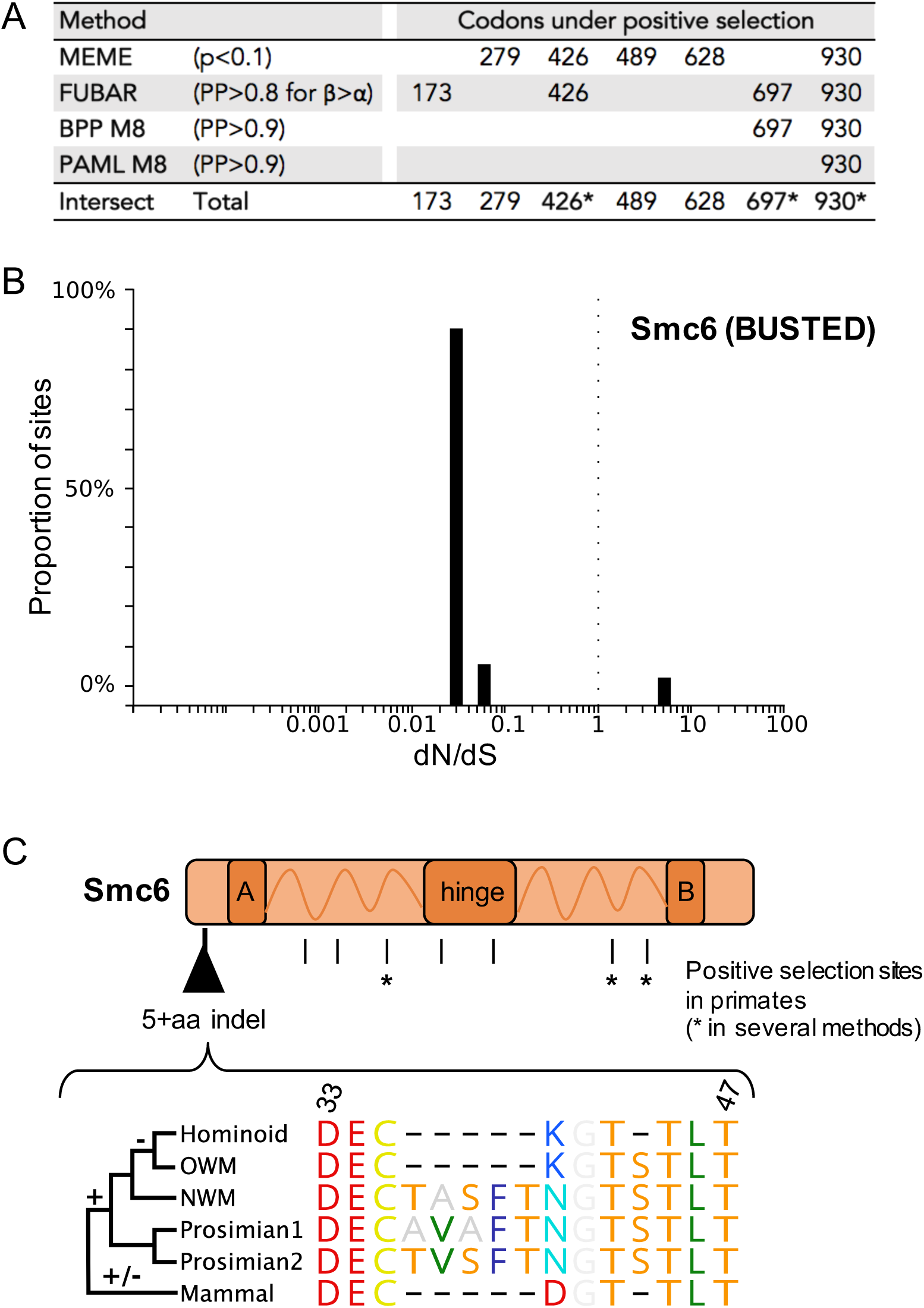
Evidence of episodic site-specific positive selection in Smc6 during primate evolution. A, Specific sites in Smc6 are under positive selection. Codon-alignments were analyzed using four different positive selection tests: MEME, which detects site-specific episodic positive selection; FUBAR, similar in a Bayesian framework; and Bio++ and PAML Codeml (M8), which detect site-specific positive selection (see Methods). The table shows the codon-sites showing significant positive selection (i.e. that passed the widely accepted p-value or posterior probability (PP) cut-offs for each method). The statistical thresholds used in each test are shown in the table. Codons identified as being under positive selection in at least two of the four tests are indicated by an asterisk. B, Graphic depicting the proportion of sites in Smc6 at a given dN/dS ratio, calculated with BUSTED. A very few number of Smc6 sites are under positive selection. Ratio dN/dS<1, negative selection; dN/dS□ =□ 1, neutrality; and dN/dS>1, positive selection. C, Marks of “genetic conflicts” in Smc6. Sites under positive selection in primates, as well as the plasticity of the N-terminal region of Smc6 in mammals are shown. Amino acid alignment was performed with MUSCLE, and residue’s color coding is from RasMol. A and B correspond to the globular domain that contain a Walker A and Walker B motif, respectively. Dashes indicate gaps. One non-primate mammal species was arbitrarily chosen for the schematic representation. The mammal sequence illustrates only one possibility of the natural inter-species sequence variations that have been important in this region (Dataset S3). For example, the extensive plasticity of the N-terminal region during bat evolution is further illustrated in Figure S2.

In addition to single codon substitutions, other forms of genetic changes may also be adaptive in a virus-host “arms-race” (21, 35, 39). In particular, recombination, gene deletion or duplication, insertions and deletions (indels) can also be advantageous for the host. These “genetic innovations” would be missed in typical methods screening for positive selection. We found some indels in genes encoding for several Smc5/6 complex subunits (Dataset S1). In particular, we found a 5-amino acid (aa) indel in the N-terminal region of Smc6 (Figure 2C). The five New World monkey species, including the two sequenced in this study, display a TASFT motif at this position. However, this 5-aa motif was not present in any of the retrieved *Catarrhini* species sequences (n=13) (Figure 2C Hominoids and Old World Monkeys; Dataset S1). To decipher if this 5-aa indel was specific to simian primates, we extended our analysis to mammals. We found a significant plasticity within this region with both indels and amino acid changes during mammalian evolution. For example, the prosimians’ proteins contain a 5-aa motif, but with amino acid differences in the TASFT motif (TVSFT and AVAFT, respectively; Figure 2C). Another remarkable example of this N-terminal plasticity was found in bats (*Chiroptera*). Most bat species contain a 5-aa stretch (residues 36-40) but the sequence differs significantly (TVSFI, TVSFT, PDPFT, TDTFT), while some bats carry an 8-10 amino acid deletion within this region (*Rhinolofus* and *Hipposideros* species) (Figure S2). Therefore, although most of the Smc6 protein sequences has been very conserved in primates and more generally in mammals, a few sites have been under significant positive selection and show substantial genetic plasticity. These signatures of genetic changes in an essential protein could be reminiscent of an evolutionary “arms-race” with pathogenic viruses.

### HBx and the woodchuck WHx counterpart promote degradation of diverse mammalian Smc5/6 complexes, independently of variations at genetic innovation sites

We next used evolutionary-guided functional assays to examine whether the marks of adaptive evolution and the identified interspecies variability of the Smc5/6 complex have functional consequences on the ability of HBx to promote its degradation. To test this, we put together a panel of mammalian cells encoding for divergent Smc5/6 (breadth of our species panel showed in Figure S4) and, importantly, with Smc6 orthologues with variations at positive selection sites and indels (Figure 3A). This panel included cells derived from various primate species, including human, tantalus African green monkey (*Chlorocebus tantalus*), vervet African green monkey (*Chlorocebus pygerythrus*), and the New World owl monkey (*Aotus trivirgatus*), as well as ferret (*Mustela putorius furo*, carnivore) and mouse (*Mus musculus*, rodent) cells. Using this panel, as well as different human cell lines, we tested the capacity of the human HBx and the woodchuck WHx counterpart to promote degradation of the heterologous Smc5/6 complex in these cells. Consistent with previous studies (Decorsiere, et al. 2016; Murphy, et al. 2016), we found that HBx transduced in various human cell lines, including HepG2 (hepatocyte carcinoma), 293T (kidney epithelial), and HeLa (epithelial adenocarcinoma), triggers a similar decrease in endogenous Smc6 and Nsmce4A protein levels (Figure S3). Similar results were obtained in U2OS (human osteosarcoma) and A549 (human lung adenocarcinoma) cell lines (data not shown). This indicates that this HBx activity is neither cell-type specific, nor affected by human polymorphism at position 697 (rs1065381).

**Figure 3.**
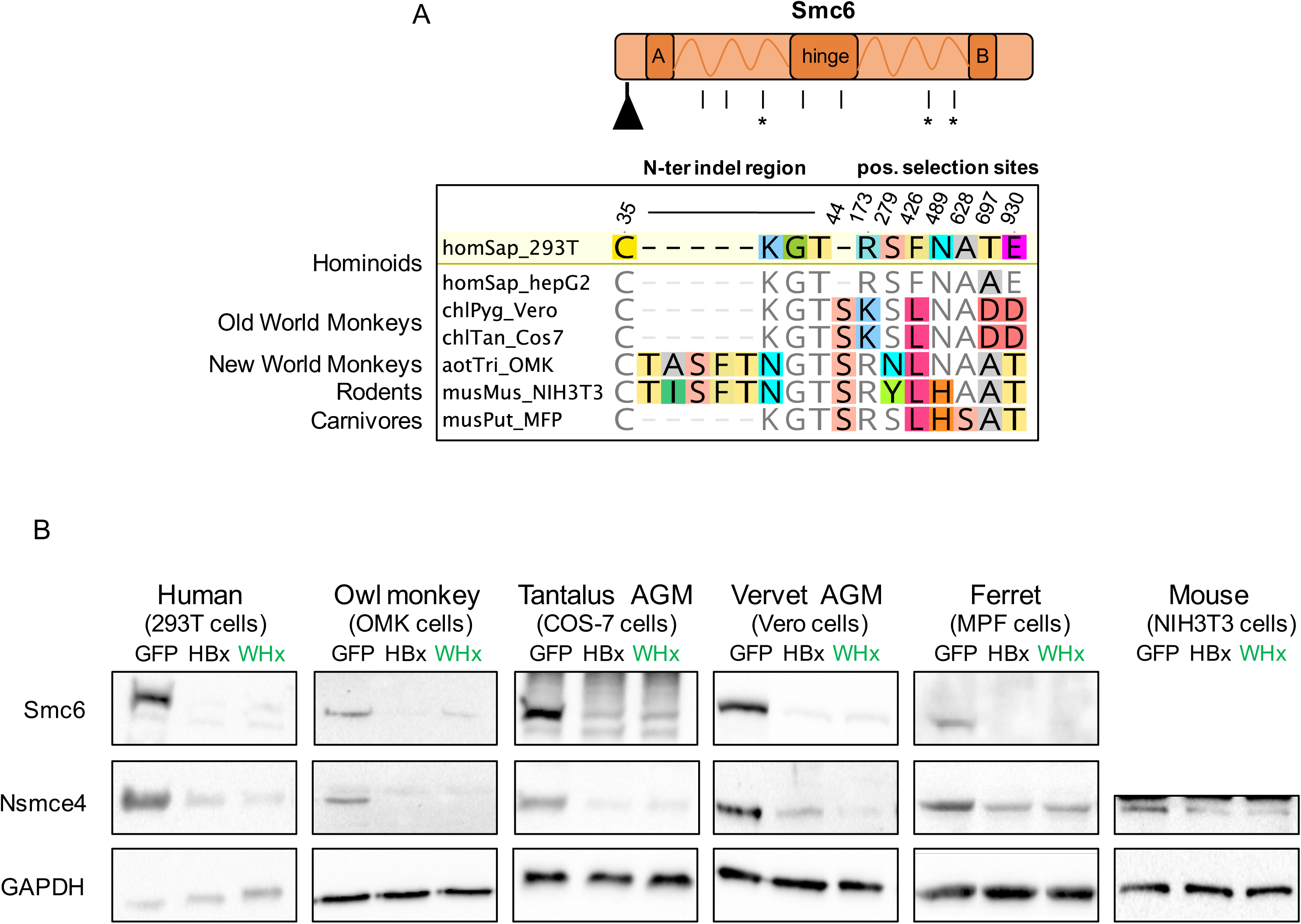
HBx and WHx can degrade the Smc5/6 complex in cells from diverse mammalian species. A, Amino acid differences at the sites of “genetic conflict” in Smc6 between the host species tested in panel B. Note that all statistically significant marks of potential evolutionary “arms-race” identified in Figure 2 are represented. Asterisks denote the sites that were found under positive selection by at least two methods. B, The human (HBx) and woodchuck (WHx) HBV X proteins promote degradation of the Smc5/6 complex in primate, rodent, and carnivore cells (n=6 species). Cells were either mock transduced or transduced with a lentivector encoding GFP, GFP-HBx or GFP-WHx. After 4-5 days, cell lysates were prepared and run on an SDS gel and probed for Smc6 and Nsmce4A, two essential subunits of the Smc5/6 complex, using primary antibodies against the human protein. Note that rodent Smc6 could not be detected. GAPDH serves as a loading control.

Then, cells from the six different mammalian species were transduced with a lentivector encoding for GFP alone, GFP-HBx or GFP-WHx. Five days later, we found that HBx and WHx expression had specifically triggered the degradation of the Smc5/6 complex in all species tested (Figure 3B). This was true even for host species like owl monkey, which harbor a 5-aa insertion in the N-terminal region of Smc6 (Figure 3A-B, Figure 2C) and for Old World monkeys that, in contrast to New World monkeys and Hominoids, are not natural hosts for hepadnaviruses (Figure 3B). Taken together, these results show that the antagonism of endogenous Smc5/6 by the viral HBx and WHx is independent of the cell type and of the mammalian host species. Overall, amino-acid differences at sites under adaptive evolution in Smc6 did not significantly impact on HBx-mediated degradation of the complex (Figure 3A-B), suggesting that HBV has not driven Smc6 adaptation in primates.

### Mammalian hepadnavirus HBx proteins show a conserved ability to counteract the restriction activity of the human Smc5/6 complex

It is unknown whether the capacity to counteract the Smc5/6 complex is an important and conserved function of mammalian hepadnaviruses that have diverged millions of years ago. To span the entire orthohepadnavirus evolutionary history, we examined, in addition to the human HBx and woodchuck WHx, the newly cloned HBx proteins from a hepatitis B virus that infects the New World wooly monkey (WMHBx) and from three viruses infecting distantly related bat species; *Hipposideros cf. ruber* (roundleaf bat), *Rhinolophus alcyone* (horseshoe bat), and *Uroderma bilobatum* (tent-making bat) (RBHBx, HBHBx, and TBHBx respectively) (Figure 4A). As shown in Figure 4B, these HBx proteins have highly divergent amino acid sequences with some regions sharing essentially no homology. Despite this high sequence and long-term divergence, all HBx proteins showed comparable ability to trigger degradation of the human Smc6 and Nsmce4A proteins, when transduced in either HepG2 or 293T cells (Figures 4C and S5).

**Figure 4.**
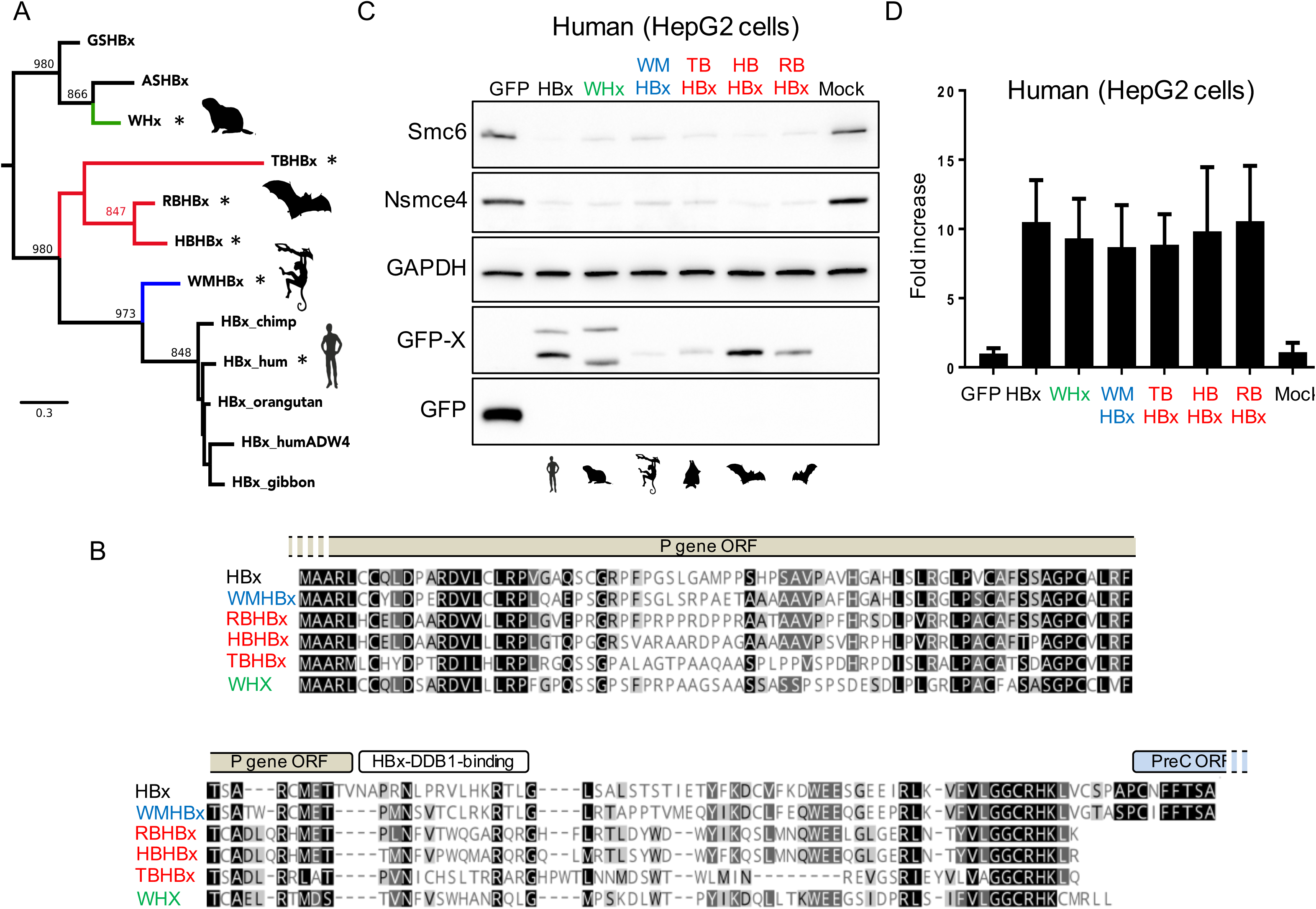
Highly divergent mammalian HBV X proteins show a conserved property of antagonizing Smc5/6 restriction in human cells. A, Phylogenetic analysis of the X proteins from hepadnaviruses that naturally infect mammals. The viral X proteins tested in our *in vitro* functional assays (C and D) are highlighted by an asterisk. Phylogenetic analysis of orthohepadnaviral X proteins was performed using a 161-amino acid alignment obtained with MUSCLE (Dataset S2) and the tree was built with PhyML and a JTT+I+G model with 1,000 bootstrap replicates. Bootstrap values (> 800/1000) are indicated at the nodes. The tree was rooted for representation purposes according to Drexler et al. 2013 (but the outgroup of orthohepadnavirus is still under debate (2)). The scale bar indicates the number of amino acid substitutions per site. We analyzed the X proteins from GSHBV, ground squirrel; ASHBV, arctic squirrel; woodchuck WHV; three bat viruses (RBHBV, HBHBV, and TBHBV naturally infecting *Hipposideros cf. ruber* (roundleaf bat), *Rhinolophus alcyone* (horseshoe bat), and *Uroderma bilobatum* (tent-making bat), respectively); wooly monkey WMHBV; human HBV and other indicated hominoids. B, Amino acid alignment of the viral X proteins used in B and C. The black-to-white gradient depicts high-to-low sequence identity. The open reading frames (ORFs) overlapping with HBx are shown, as well as the DDB1-binding region in the human HBx protein (70). Of note, we suspect that all orthohepadnavirus X proteins are able to recruit the DDB1 complex although the domain by which they do so appears non-homologous at the sequence level. C, Degradation of the human Smc5/6 complex by mammalian X proteins. Human hepatoma HepG2 cells were either mock transduced (Mock) or transduced with a lentivector expressing GFP alone or fused to the indicated X proteins. Western blot analysis of the endogenous Smc6 and Nsmce4A was performed (see Methods). GAPDH serves as a loading control. Supplementary Data are found in associated Figure S5. D, Effect of mammalian X proteins on transiently transfected reporter gene activity. HepG2 cells were transfected with a luciferase reporter construct and the next day transduced with lentiviral vectors expressing the indicated proteins as above. At day 5-7, the luciferase activity was measured and the fold-increase of RLU (relative light units) versus the GFP control condition (set at 1) is shown. The mean of two independent experiments is shown, along with the standard deviation.

Previous work has shown that the Smc5/6 complex binds episomal DNA templates to block transcription and that inactivation of the complex leads to an increased episomal gene expression (Decorsière et al., 2016). Accordingly, a similar increase in expression of a transiently transfected episomal luciferase reporter construct was observed in all orthohepadnavirus HBx protein-expressing cells (Figure 4D). Thus, the ability to degrade and to counteract the restriction activity of the Smc5/6 complex is conserved among mammalian hepadnavirus HBx proteins.

### Divergent mammalian HBx proteins efficiently rescue replication of a human HBx-deficient hepatitis B virus in primary human hepatocytes

We finally tested whether the HBx proteins from non-human orthohepadnaviruses would substitute for human HBx in an HBV replication assay. Primary human hepatocytes (PHHs) were transduced with lentiviruses expressing GFP alone or fused to one of the six orthohepadnavirus HBx proteins, and four days later infected with the wild-type HBV or an HBx-deficient HBV mutant (HBVAX) (Figure 5A). As shown previously, human HBx provided *in trans* fully rescued the replication defect of HBV∆X as measured by HBe and HBs antigen secretion (Decorsière et al 2016). Strikingly, the HBx proteins from wooly monkey (WMHBx), woodchuck (WHx), and the three bat (TBHBx, HBHBx, and RBHBx) viruses were all capable of restoring HBV∆X replication to comparable levels as shown by HBe and HBs antigen secretion (Figure 5A). This occurred in the absence of changes in viral cccDNA levels (Figure 5B) and was accompanied by a concomitant decrease in Smc6 protein levels (Figure 5C). These results provide functional evidence that orthohepadnavirus HBx proteins have a conserved capacity to antagonize the antiviral function of Smc5/6 and that this occurs in a species-independent fashion.

**Figure 5.**
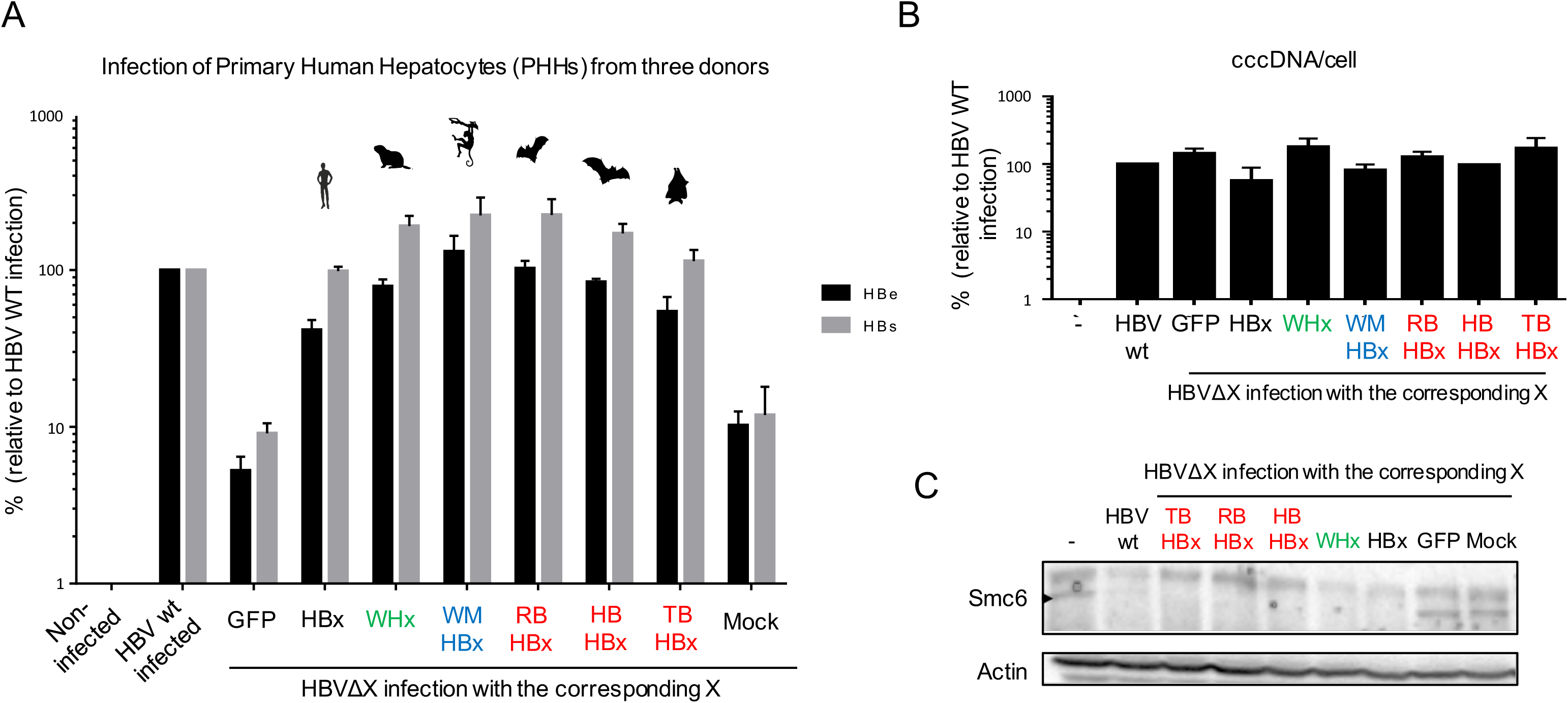
The X proteins from six orthohepadnaviruses can all fully rescue the replication defect of an HBx-defîcient HBV in primary human hepatocytes (PHHs) A, PHHs were mock-transduced or transduced with GFP or the indicated X proteins and infected with wild-type HBV or an HBx-deficient HBV (HBV ΔX). HBe and HBs antigen secretion was quantified 7 days later by ELISA. Antigen concentrations are relative to wildtype HBV, which was set to 100. Data are mean ± SEM of independent experiments performed with three different PHH donors. B, The HBV cccDNA levels were measured at day 7 post-infection by real-time PCR. Values are expressed relative to β-globin mRNA levels to normalize to cell number. The results are the mean +/-SEM for the levels seen in PHHs from two donors. C, Smc6 degradation in PHHs expressing X proteins from different HBV lineages. Protein extracts where prepared from the cells above and Smc6 protein levels were analyzed by Western blot. Actin serves as a loading control. Supplementary Data are found in associated Figure S6.

## Discussion

In this study, we found that the HBx proteins from mammalian hepadnaviruses that have diverged millions of years ago and have very little sequence homology all retain the capacity to degrade the Smc5/6 complex and to counteract its antiviral activity in a species-independent manner. The antiviral function of the Smc5/6 complex against hepadnaviruses has therefore been an important immune defense mechanism in mammals.

We traced the evolutionary history of the genes encoding the six components of the Smc5/6 complex, as well as the genes for the Smc1-4 core subunits of cohesin and condensin. We show that Smc1-4 and the Smc5/6 complex have been highly conserved in primates. The only exception is the *Smc6* gene, which shows some signatures of adaptive evolution in primates and mammals. These include several sites under positive selection and insertion/deletion events during mammalian evolution, which could be reminiscent of a virus-host evolutionary “arms-race” To examine if HBV, which is currently the only virus reported to be restricted by the Smc5/6 complex, has contributed to shape the evolution of the complex, we tested several orthohepadnavirus HBx proteins for their ability to antagonize the Smc5/6 complex across species. We found that all the HBx proteins we tested, including those encoded by distantly related bat HBVs, were equally efficient in antagonizing Smc5/6 in human and other mammalian cells. This conserved property suggests that the interaction between the Smc5/6 complex and HBx is independent of the sites under adaptive evolution that we identified in this study. Thus, HBx does not seem to have driven the evolution of Smc5/6 in primates and, more broadly, in mammals.

Overall, we did not identify strong signatures of positive selection in the Smc5/6 complex as typically found in other known restriction factors (18, 21, 39, 40). This is surprising given that the antagonistic relationship between HBx and Smc5/6 has been conserved in orthohepadnaviruses (this study) and has likely been played out over tens of millions of years (1). There are at least three possible explanations for this apparent inconsistency. Firstly, Smc5/6 has an evolutionarily highly conserved architecture and it performs fundamental functions in cellular genome maintenance, and thus has likely evolved under strong purifying selection. This is in contrast to most other described restriction factors, such as APOBEC3G and TRIM5, that are dedicated to their antiviral intrinsic immune functions (40, 41). However, our findings show similarities both at the evolutionary and functional levels with the Serine incorporator proteins SERINC3 and SERINC5. These cellular proteins were recently found to act as restriction factors against lentiviruses (42-45). Despite their antiviral function and their antagonism by the lentiviral Nef protein, SERINC3 and SERINC5 also show little evidence for positive selection (Murrell, et al. 2016). Like Smc5/6, the SERINC proteins have important cellular functions and are not part of the interferon signaling pathway. The Smc5/6 complex and the SERINC proteins may therefore fall into a separate category of restriction factors that have dual antiviral and essential cellular functions, and therefore have limited evolutionary opportunities. In an extreme scenario, host proteins might not be able to “escape” from the viral antagonist and it is likely that viruses have adapted to target such constrained host proteins (i.e. “viral strategy”) (36, 46). The thorough evolutionary analysis of such constrained restriction factors would benefit from novel bioinformatic methodologies specifically designed to identify adaptive evolution that operates on a background of strong purifying selection and that may further involve genetic innovations other than site-specific positive selection (e.g. indels, recombination).

A second possible explanation for the lack of strong positive selection in the Smc5/6 complex is that orthohepadnaviruses may not have been strong drivers of mammalian genome adaptation. Indeed, acute HBV infection is mostly asymptomatic, and when evolving into chronic infection the symptoms and associated morbidity appear late in life, after the reproductive age (WHO IARC Monograph). Arguing against this interpretation are the findings of Enard and colleagues, who provided evidence that most cellular genes reported in the literature to encode HBV-interacting proteins show a strong excess of adaptation (36). This, however, remains to be functionally demonstrated because, to our knowledge, no study to date has reported evidence for a direct evolutionary “arms-race” between an HBV protein and a cellular factor.

A third possibility is that HBx interacts with the Smc5/6 complex indirectly, through a yet unknown intermediate cellular protein. Although we cannot formally exclude this possibility, none of our studies so far suggests that this is the case. In particular, no common cellular protein in addition to the DDB1 subunit of the E3 ligase and the six Smc5/6 subunits was recovered from HBx-expressing cells in pull down experiments with either HBx or Smc5/6, as would be expected for an adaptor protein bridging HBx to the complex (FA and MS, unpublished). We can further exclude any “evolutionary arms-race” between DDB1 and HBV, because DDB1 is under strong purifying selection (data not shown) and our assays in heterologous species cells allow us to robustly exclude any species-specificity between HBx and a hypothetical endogenous intermediate factor. Nevertheless, the exact HBx-Smc5/6 interface remains unknown. Evolutionary and mutagenesis analyses of HBx did not allow us to identify the viral determinants of the interaction with Smc5/6 (data not shown). The analysis of structural/docking models ((47, 48) for example) might contribute to solve this virus-host interface.

It is really remarkable that all the orthohepadnavirus HBx proteins that we tested antagonize the Smc5/6 complex and share the ability to fully substitute for human HBx in an HBV infection assay using primary human hepatocytes, especially given their low sequence identity (<38% identity for some pairs) and their time of divergence. These findings suggest that the Smc5/6 antiviral restriction activity is conserved and has been essential among mammals and provide additional evidence that antagonizing the Smc5/6 complex is a major function of HBx during HBV infection (49-51). Our findings further imply that, in contrast to what has been documented for other restriction factors (52), Smc5/6 does not act as a species-barrier to the potential zoonotic transmission of bat or primate HBVs to humans (49-51, 53).

HBVs are however not restricted to mammals as they are also found in birds, fishes, and amphibians (2, 54, 55). Because these latter viruses appear to lack an *HBx* gene (1, 2, 54), it would be interesting to explore if they are also restricted by Smc5/6, and if so, what strategy do these divergent hepadnaviruses use to circumvent the antiviral restriction.

Finally, because mammalian *Smc6* has some evidence of adaptive evolution independently of HBV pressure, it raises the possibility that the *Smc6* gene has been engaged in an evolutionary “arms-race” with other pathogenic viruses. For example, the evolutionary adaptive changes identified in the lentiviral restriction factor MxB have not been driven by lentiviruses but likely by other pathogens (44, 56). In addition, restriction factors with a broad antiviral spectrum, such as MxA or PKR, have been evolutionarily driven by several pathogens (57, 58). It will be interesting to determine whether other DNA viruses are restricted by the Smc5/6 complex and, if so, whether they have contributed to shape the evolution of mammalian Smc5/6 complex.

## Methods

### Cell lines and culture

Cells were maintained in DMEM supplemented with 10% fetal calf serum. Human cell lines used in the study were: human embryonic kidney 293T cells and HeLa cells (gifts from Andrea Cimarelli, CIRI Lyon), as well as the human hepatoma cell lines HepG2 (ATCC HB-8065). Old World monkey cells used were: tantalus African green monkey AGM (*Chlorocebus tantalus*) COS-7 cells (a gift from Branka Horvat, CIRI Lyon) and vervet AGM (*Chlorocebus pygerythrus*) Vero cells (a gift from Andrea Cimarelli, CIRI Lyon). New World monkey cells were: Owl monkey (*Aotus trivirgatus*) OMK cells (CelluloNet Lyon) and Coton-headed tamarin (*Saguinus oedipus*) B95a cells (a gift from Branka Horvat). We also used ferret (MPF, *Mustela putorius furo*, from Branka Horvat) and mouse cells (NIH3T3, a gift from Theophile Olmann’s lab, CIRI Lyon). The species identity of all the cell lines used in this study was confirmed by amplification and sequencing of the Cytochrome b and/or beta-actin (data not shown).

### Expression Plasmids

The lentivirus vectors pWPT expressing GFP, GFP-HBx, and GFP-WHx have been previously described (59-61). The X coding regions (synthesized by Genewiz) from hepadnaviruses infecting the New World wooly monkey (*Lagothrix*) (WMHBx), three distant bat species including the roundleaf bat (*Hipposideros cf. ruber*), the horseshoe bat (*Rhinolophus Alcyone*), and the tent-making bat (*Uroderma bilobatum*) (RBHBx, HBHBx, TBHBx, respectively) (51) were expressed from the same vector. The X insert was ligated to the pWPT-GFP backbone between the PstI and NotI sites following the T4 DNA (NEB-M0202) ligation protocol. The plasmid was then transformed following the High Efficiency Transformation Protocol using NEB 10-beta Competent E. coli (NEB-C3019). All the X-expressing plasmids were checked by Sanger sequencing using eGFP-F primer, 5’-CAT GGT CCT GCT GGA GTT CGT G-3’, and pWPT-R, 5’-CAT GGT CCT GCT GGA GTT CGT G-3’.

### Transfection and transduction

For recombinant lentivirus production, plasmids were transfected in 293T cells by the Calcium Phosphate-method. Briefly, 4.5×10^6^ cells were plated in a 10cm-dish and transfected 12h later with 15 μg of the X protein-expressing lentiviral vectors, 10 μg of packaging plasmid (psPAX2, gift from Didier Trono (Addgene plasmid # 12260)), and 5 μg of envelope (pMD2G, gift from Didier Trono (Addgene plasmid # 12259)). Media was changed 6h posttransfection. After 48h, the viral supernatants were collected and stored at −80°C. The viral supernatants were titrated by transducing 293T or HeLa cells and measuring the GFP expression by FACS 5 days later. The different mammalian cell lines were transduced by plating 0.1-0.5 ×10^6^ cells in 12-well plates and adding 4-6h later appropriate volumes of viral supernatants to obtain at least 75% of GFP positive cells. Because of the strong post-entry block against the lentiviral vector in OMK cells due to the owl monkey TRIMcyp (62), cyclosporine A was added (final concentration 2.5μM) to increase transduction efficiencies in these cells. Five days post-transduction, cells were collected for FACS and Western blot analysis. For AGM Cos-7 cells, transduction efficiencies were low and GFP-positive cells were isolated using FACS AsriaII sorting.

### Luciferase reporter assay

Transfection of plasmid DNA in HepG2 cells was performed using X-tremeGENE HP DNA Transfection Reagent (Roche) following the manufacturer’s instructions. For luciferase reporter gene assay cells were typically seeded at a density of about 6×10^5^ cells per 30mm diameter well (1×10^5^ cells/cm2) and reverse-transfected with 30 ng of reporter plasmid DNA pCMV-Gluc (New England Biolabs) and 2 μg of empty EBS-PL vector. The next day cells were transduced with lentiviral vectors encoding GFP or the diverse GFP-tagged viral X proteins. Five to seven days later GLuc activity was measured by adding 5 μL of the cell supernatant sample to 50 μL room temperature assay buffer (100 mM NaCl, 35 mM EDTA, 0. 1% tween 20, 300 mM Sodium ascorbate, 0,8 μM coelenterazine in PBS 1X), and immediately measuring luminescence in a luminometer (Glomax, Promega).

### Western blotting

Western blot analyses were performed as previously described (7). Cells were disrupted in 2% SDS and briefly sonicated. The protein concentration was estimated and normalized using the BCA Protein Assay (Novagen). The membranes were probed with 1:5,000 mouse monoclonal anti-GFP antibody (Roche 11814460001) to detect the GFP-tagged X proteins, 1:1,000 mouse monoclonal antibodies against Smc6 (Abgent AT3956a), 1:1,000 rabbit polyclonal anti-Nse4 (Abgent AP9909A), 1:10,000 mouse monoclonal anti-GAPDH (Sigma-Aldrich G8795) antibodies. Horseradish peroxidase-conjugated sheep antirabbit or anti-mouse IgG (Amersham Biosciences, at 1:5,000) were used as secondary antibodies. Detection was performed by ECL (Pierce).

### PHH isolation, HBV infection, ELISA

PHHs were isolated from human liver tissue as previously described (63) from three donors. They were infected with PCR normalized HBV or HBVΔX at an MOI of 500 viral genome equivalents per cell (6). An ELISA (Autobio Diagnostics) was used to determine the amount of HBe and HBs antigens in culture media from infected or mock infected PHHs. The efficiency of infection was controlled by quantitative PCR analyses specific for the viral cccDNA using beta-globin levels to normalize cell numbers. DNA from infected and transduced cells was extracted using the Epicentre kit. For the cccDNA analysis, T5 exonuclease (TSE44111K Epicentre) was added and the DNA further incubated at 37°C for 30 min and 30 min at 70°C. Specific TaqMan probes and primers were then used to assess the cccDNA content as described in (64).

Transduction levels were estimated by the GFP expression in transduced cells by FACS analyses (BD FACS Calibur).

### *De novo* sequencing of Smc5/6 genes

Total RNA was extracted from 10^7^ cells following the manufacturer instructions (NucleoSpin RNA Blood, Macheray-Nagel 740200). Reverse transcription was performed using the SuperScript III Reverse Transcriptase (ThermoFisher 18080) with random hexamers and oligo(dT). Single-round PCR of overlapping fragments were generated using the Q5 high fidelity DNA polymerase (NEB M0491) following the manufacturer instructions. Sequences of the *Smc6* gene from owl monkey (OMK), coton-headed monkey (B95a), tantalus and vervet AGM (Cos7 and Vero), human (293T and HepG2), and ferret (MPF) cells were obtained. Primers used for the amplification and the sequencing are given in Table S1.

### Host phylogenetic analyses

Sequences of primate or mammalian orthologous genes were retrieved from publically available databases using UCSC Blat and NCBI Blastn with the human sequence as the query. After including sequences obtained from *de novo* sequencing in this study, the sequences of the orthologues were codon-aligned using Muscle (65) with minor adjustments (alignments are in Dataset S1). The name of each sequence has been uniformed: the first three letters of the host genus name followed by the first three letters of the species name (e.g. papAnu for *Papio Anubis*, or macMul for *Macaca mulata*). When sequences were retrieved using Blat on the primate full-genome assembly, the name of the genome assembly was used (e.g. panTro4 for *Pan troglodytes,* or hg38 for human). The synteny of each locus of interest was analyzed in UCSC using Blat and the Genome Browser. When necessary, the genomic sequences were further retrieved and aligned to a reference gene to determine the pseudogenes and the gene orders (Figure S1).

We used GARD from HYPHY to perform the recombination analyses with a cut-off at p<0.05 (27, 66). PhyML was used for the phylogenetic reconstructions with a HKY+G+I model and aLRT or 1,000 bootstrap replicates for branch support (67).

### Positive selection analyses

Maximum-likelihood tests to assay for positive selection were performed using three platforms: HYPHY from Kosakovsky Pond and colleagues, PAML from Yang and colleagues, and Bio++ from Dutheil, Guéguen and colleagues (30, 31, 66). In HYPHY, we used PARRIS to detect if a subset of sites in the alignment have evolved under positive selection (29). We further used the more recent BUSTED method, which detects gene-wide evidence of positive selection within a codon-alignment (28). In PAML, we used Codeml with the corresponding gene tree inferred with PhyML as input. Parameters were checked using M0 model. The gene-coding sequence alignments were fit to models that disallow (M7) or allow (M8) positive selection. For several genes, we did not get convergence certainly because genes were too conserved and the sum of dS across the tree was too low. We therefore exclude the Codeml and Bio++ analyses of seven genes, for which the p or the q parameters of the beta distribution was extreme (<0.05 or >99 (30)) (Figure 1C, NA). The likelihood of models was compared using a chi-squared test to derive p values.

For the *Smc6* gene, for which we found evidence of positive selection in some of the previously described methods, we further analyzed (i) which lineage(s) during primate evolution have been subjected to positive selection (i.e. when did the gene experience rapid evolution?) and (ii) which sites have been under positive selection (i.e. where has the gene evolved more rapidly than expected?). For branch-specific analyses, we used BS-REL from HYPHY, which identifies if certain lineages have undergone positive selection (38). To detect episodic site-specific positively selected sites, we used MEME from HYPHY (68). We also ran FUBAR in Datamonkey, which uses Bayesian inference to detect positive and negative selection at individual sites (69). In Bio++ and Codeml, we used the Bayesian Posterior Probability and the BEB analysis from the respective M8 model to identify codons with dN/dS>1 (sites with posterior probability > 0.90 are presented here, Figure 2A).

### Virus phylogenetics

The nucleotide sequences of the *X* gene from orthohepadnaviruses were retrieved using Blastn with the human HBx as the query. Only one sequence per orthohepadnaviral lineage was retained for this analysis. We performed the amino acid alignment with Muscle (total length, 161aa) and we used PhyML for the phylogenetic reconstructions with a JTT+G+I model and 1,000 bootstrap replicates for branch support (67).

### Ethics statement

PHHs were prepared from adult surgical liver resections provided by Pr Michel Rivoire’s, Pr. Jean-Yves Mabrut’s and Dr. Guillaume Passot’s departments. Approval from the local and national ethics committees (French Ministry of Research and Education numbers AC-2013-1871, DC-2013-1870, DC-2008-235) and written informed consent from patients were obtained.

### Accession numbers

All the new *Smc6* sequences are available at GenBank under accession numbers MF624755 to MF624761.

## Acknowledgements

We would like to very sincerely thank Léa Picard and Laurent Guéguen for the Bio++ analyses and input on evolutionary analyses, Véronique Barateau for technical support with the flow cytometry analyses, Andrea Cimarelli, Olivier Hantz and Isabelle Chemin for scientific discussions, and Loic Peyrot, Anaelle Dubois, and Francoise Berby for technical assistance. We thank Maud Michelet, Jennifer Molle, Loic Peyrot, Anaelle Dubois, Océane Floriot, Laura Dimer, Marie-Laure Plissonnier and Julie Lucifora (CRCL) for the isolation of primary human hepatocytes, and Pr Michel Rivoire’s, Pr. Jean-Yves Mabrut’s and Dr. Guillaume Passot’s staff for providing liver resections. We thank Michael Emerman, Andrea Cimarelli and Oliver Fregoso for their comments on the manuscript. We thank all the contributors of publically available genome sequences. This work was supported by an amfAR Mathilde Krim Phase II Fellowship (#109140-58-RKHF to LE), an FRM *“Projet Innovant”* grant (#ING20160435028 to LE), a FINOVI “recently settled scientist” grant (to LE), an ANR LabEx ECOFECT grant (GrASP to LE), an ANRS grant (#ECTZ19143 to LE), a fellowship from La Ligue Contre le Cancer (to LG), a grant from the Swiss National Science Foundation (310030-149626 to MS and FA) and by the Canton of Geneva.

## Author contributions

Conceptualization: FA, LG, MS, LE

Formal analysis: FF, FA, LG, MS, LE

Funding acquisition: JH, MS, LE

Investigation: FF, FA, LG, AP, JH, MS, LE

Methodology: FA, LG, LE

Project administration: LE

Resources: FF, FA, LG, JH, MS

Supervision: FA, JH, MS, LE

Writing – original draft: FF, LE

Writing – review and editing: FF, FA, LG, JH, MS, LE

## Supplementary Figures

**Figure S1.**
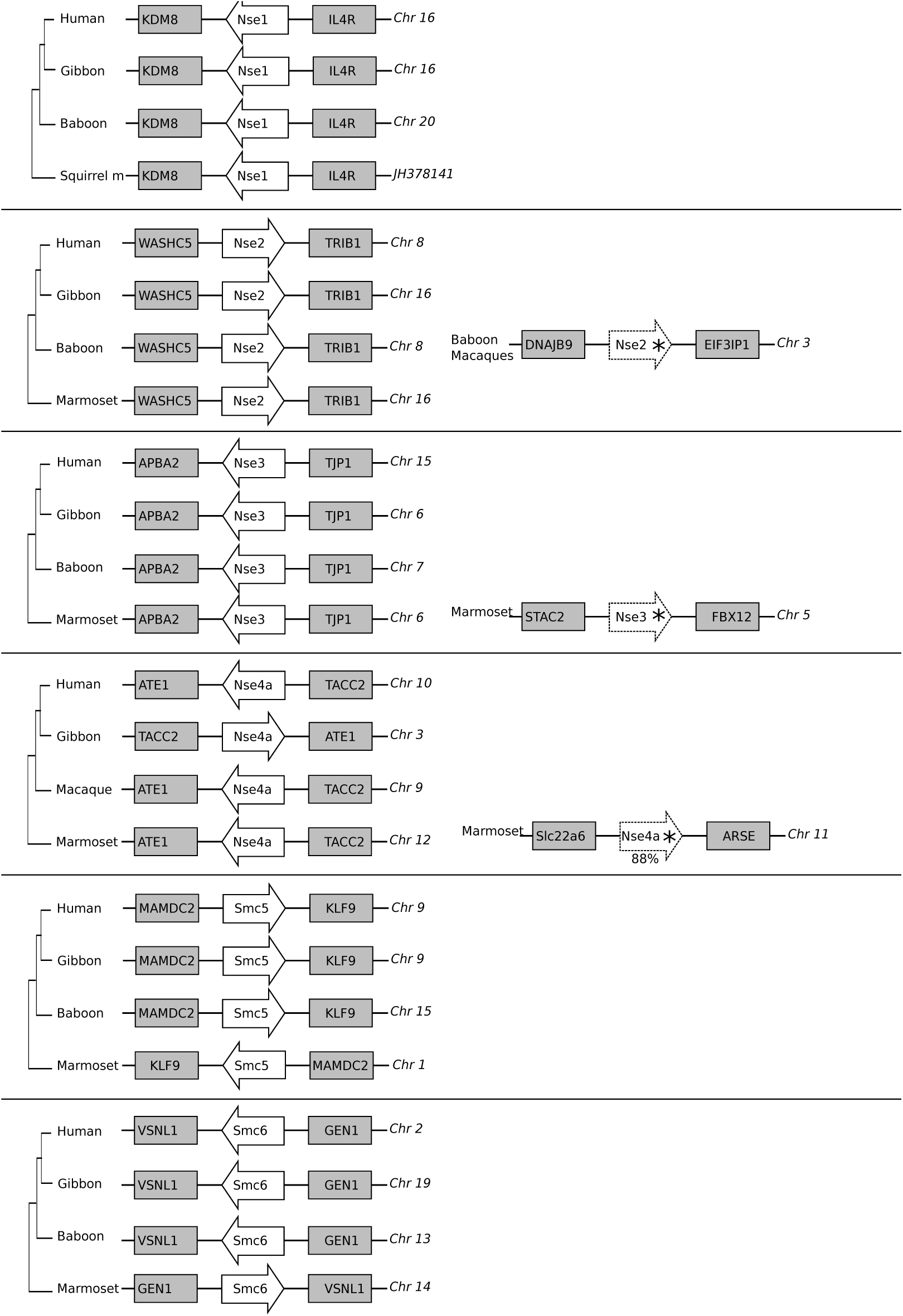
Synteny conservation of Smc5/6 complex genes during primate evolution. The six essential genes of the complex were analyzed in UCSC with the Genome Browser and Blat. For each gene, at least four primate species were fully analyzed: two great apes (human and gibbon), at least one Old World monkey (baboon and/or macaque), and at least one New World monkey (squirrel monkey and/or marmoset). The genes of interest are shown with their orientation (white arrow). Nscme1-4 are abbreviated Nse1-4. The genes in the flanking regions are shown in grey boxes. The reference of the chromosome or assembly is given on the right. For three subunits (Nscme2, Nsmce3, and Nscme4a), we found evidence of potential pseudogenes with premature stop codons (annotated with an asterisk) in non-human primate species.

**Figure S2.**
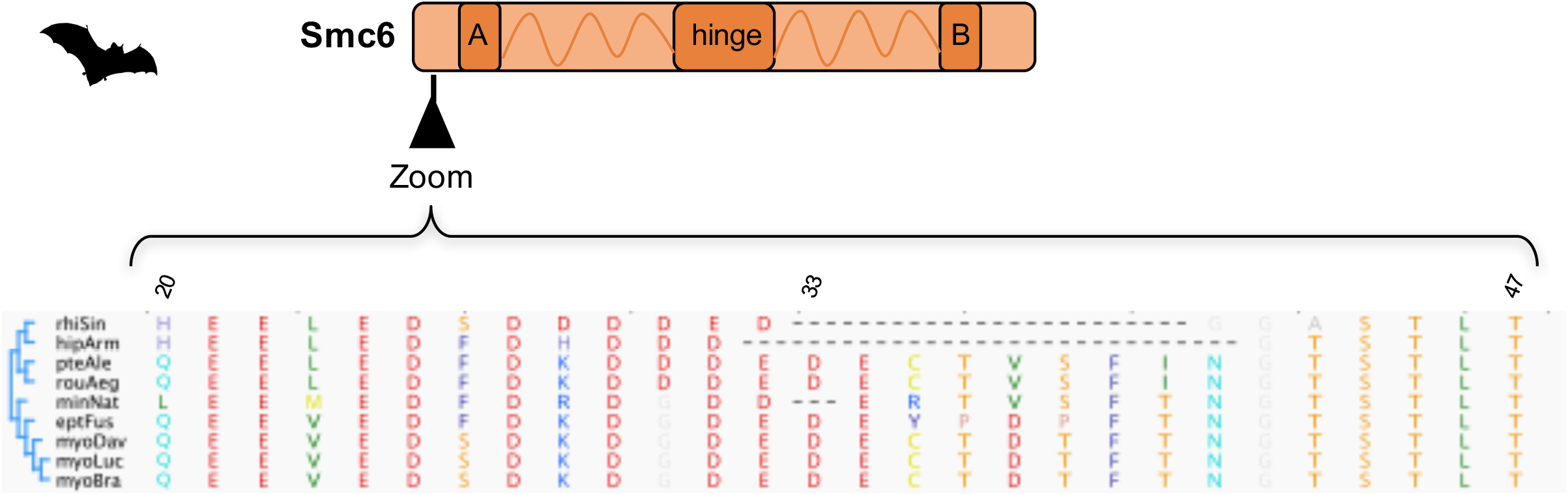
Plasticity of the N-terminal region of Smc6 during the bat’s evolution. Truncated amino acid alignment (region 20-47) of Smc6 sequences from bats (names of bat species are abbreviated: three first letters of the genus followed by the three first letters of the species). On the left, cladogram of the bat Smc6. Amino acid alignment was performed with Muscle, and residue’s color coding is from RasMol (in Geneious). Dashes indicate gaps.

**Figure S3.**
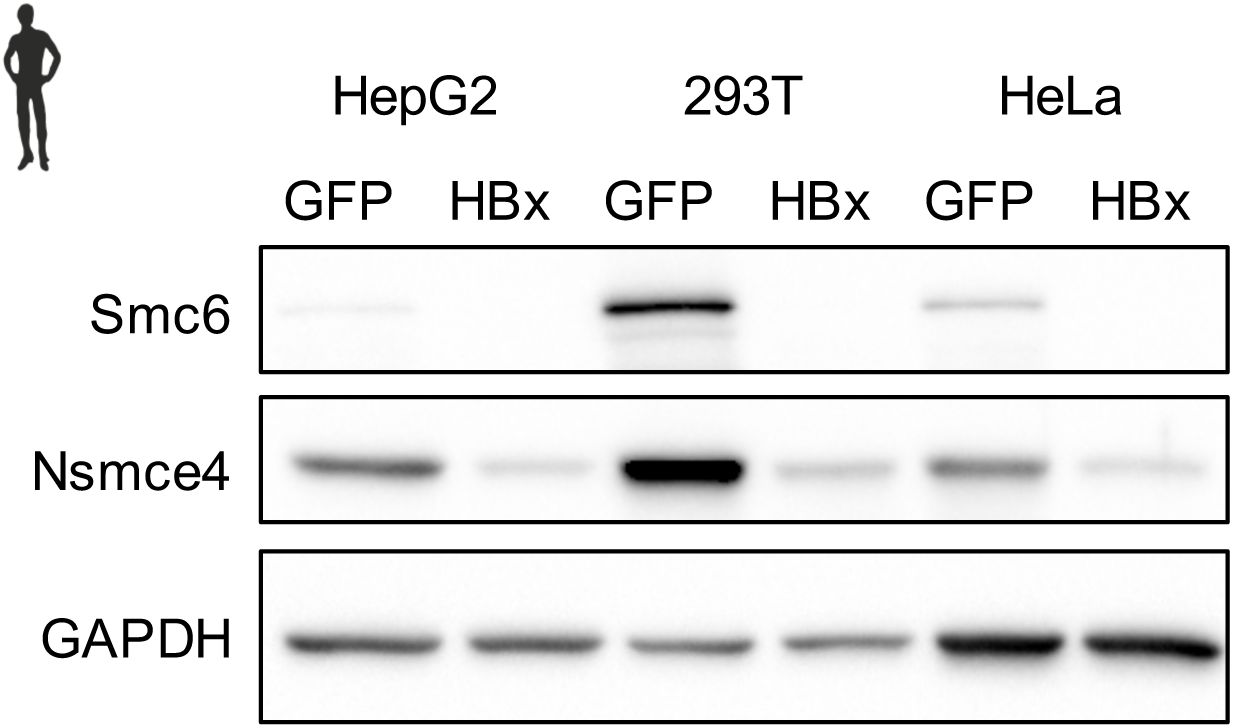
HBx from human HBV degrades the human Smc5/6 complex in a cell-type independent manner. Protein expression of endogenous Smc6 and Nscme4A from three human cell lines (HepG2, 293T, HeLa cells) that were previously transduced with a lentivector expressing GFP only (GFP) or the HBx protein from human HBV with the GFP reporter (HBx). GAPDH serves as a loading control.

**Figure S4.**
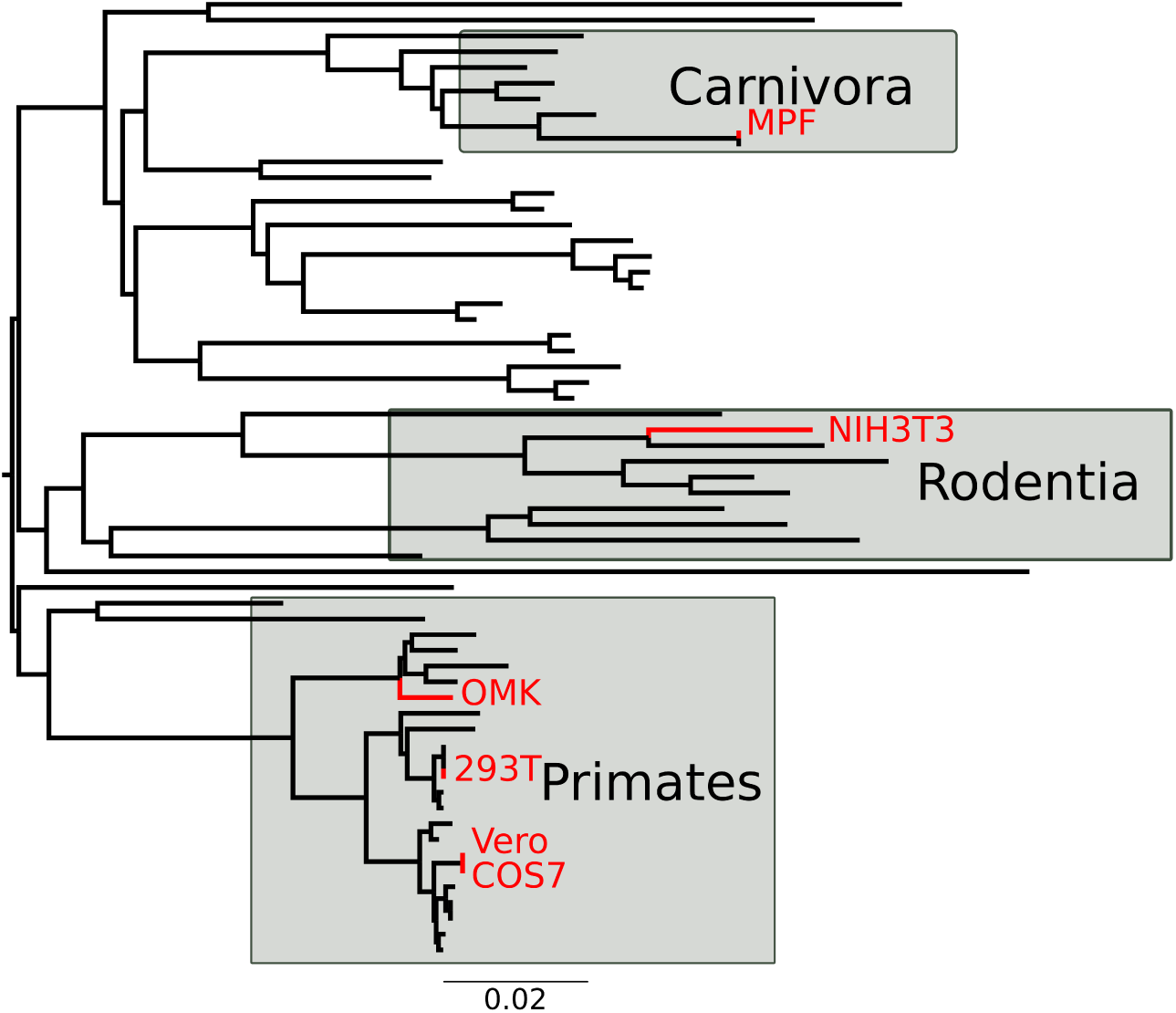
Phylogeny of the mammalian *Smc6* highlighting the species that were functionally tested in Figure 3. Our panel of host species cell sample Smc6 diversity. Nucleotide sequences were aligned with MUSCLE and phylogeny was built with PhyML and a HKY+I+G model with aLRT as statistical support (Dataset S5, Table S2). The tree was rooted according to the accepted mammalian species tree. Host species that were functionally tested are highlighted with the name of the corresponding cell line in red.

**Figure S5.**
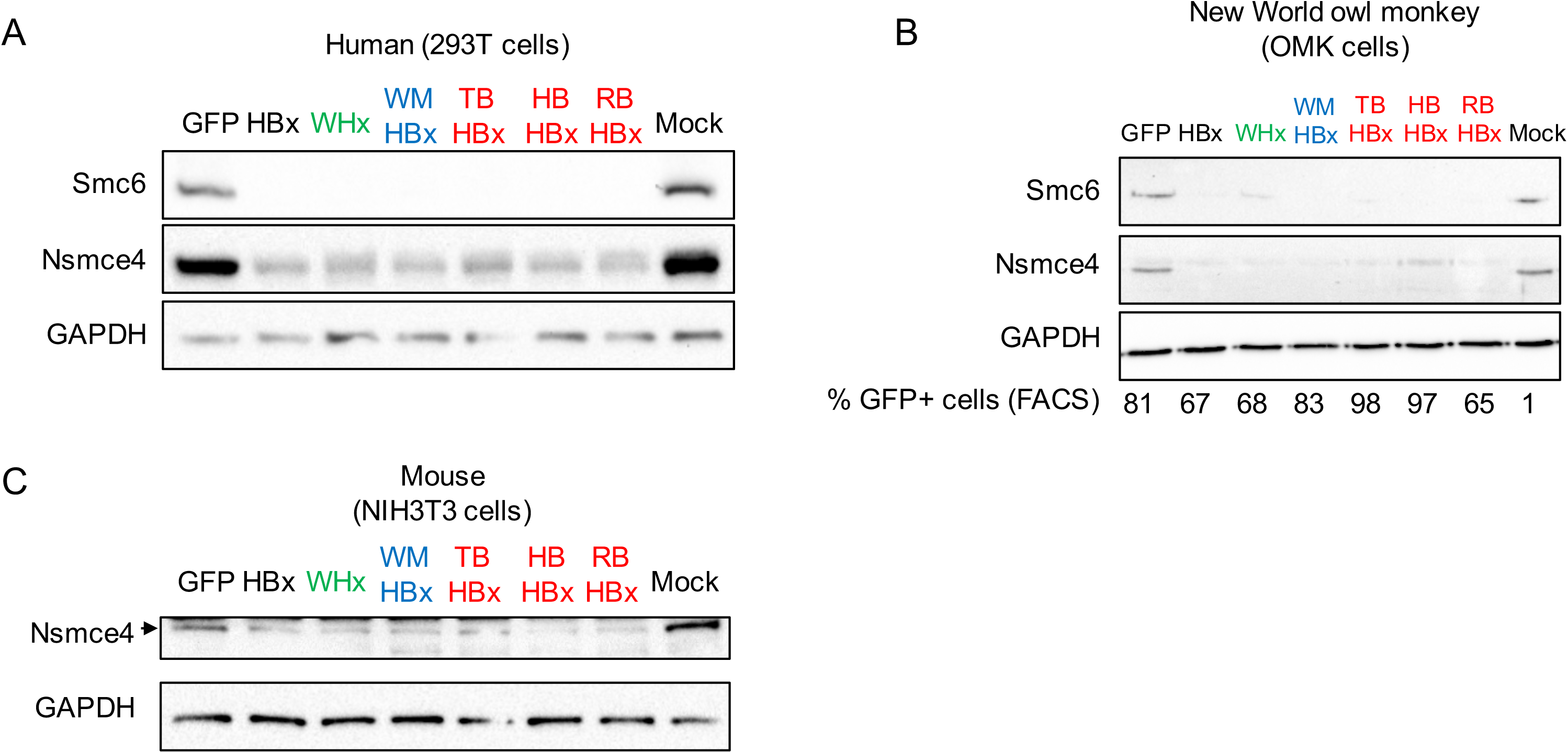
Associated to Figure 4. The X proteins from six divergent mammalian hepadnaviruses degrade the Smc5/6 complex from (A) human (293T cells), (B) New World owl monkey (OMK cells), and (C) mouse (NIH3T3 cells). As in Figure 4, human 293T cells (A), owl monkey OMK cells (B), or mouse NIH3T3 cells were transduced with a lentivector expressing only GFP (control), or the GFP-fused X protein from diverse hepadnaviruses: human HBV, woodchuck WHV, wooly monkey WMHBV, and viruses from three bat species (RBHBV, HBHBV, and TBHBV naturally infecting *Hipposideros cf. ruber* (roundleaf bat), *Rhinolophus alcyone* (horseshoe bat), and *Uroderma bilobatum* (tent-making bat), respectively), or a mock control. Western blot analysis of the endogenous Smc6 and Nsmce4A was performed (see Methods). Note that mouse Smc6 could not be detected. GAPDH serves as a loading control.

**Figure S6.**
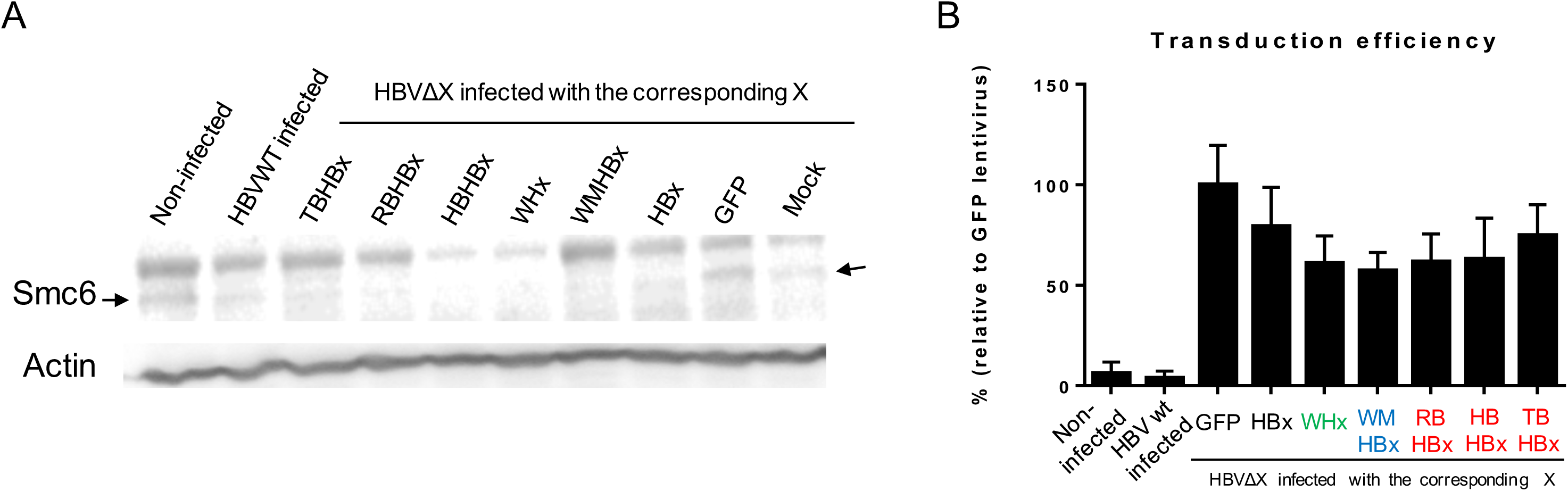
Associated to Figure 5. The X proteins from six orthohepadnaviral lineages can all rescue the replication of an HBx-deficient HBV in primary human hepatocytes (PHHs). A, Smc6 degradation was seen in PHHs expressing the X protein from different lineages. Protein extracts where prepared from infected cells on day 7 and Smc6 levels analyzed by western blot. Data shown is a representative gel from the PHHs from one donor (different from Figure 5C). B, Transduction levels were estimated by the GFP expression (from HBx-GFP or GFP alone) in transduced cells by FACS analyses (BD FACS Calibur).

## Supplementary Tables

**Table S1.** Primers used to amplify and sequence endogenous Smc6 from different mammalian cell lines.

**Table S2.** Species used for the phylogeny of the mammalian *Smc6* (Figure S4). In red: the genes that were newly sequenced

## Supplementary Dataset

**Dataset S1.** Host gene alignments used in the study (fasta files) and phylogenetic analyses (newick format)

**Dataset S2.** Orthohepadnaviral HBx amino acid alignment (interleaved phylip format) **Dataset S3.** Phylogenetic analyses of Smc5/6 in mammals (fasta and newick format)?

